# Diffusion property and functional connectivity of superior longitudinal fasciculus underpin human metacognition

**DOI:** 10.1101/2020.03.17.994574

**Authors:** Yunxuan Zheng, Danni Wang, Qun Ye, Futing Zou, Yao Li, Sze Chai Kwok

**Affiliations:** Shanghai Key Laboratory of Brain Functional Genomics, Key Laboratory of Brain Functional Genomics Ministry of Education, Shanghai Key Laboratory of Magnetic Resonance, Affiliated Mental Health Center (ECNU), School of Psychology and Cognitive Science, East China Normal University, Shanghai, China; Division of Natural and Applied Sciences, Duke Kunshan University, Kunshan, Jiangsu, China; NYU-ECNU Institute of Brain and Cognitive Science at NYU Shanghai, Shanghai, China; Shanghai Changning Mental Health Center; School of Biomedical Engineering, Shanghai Jiao Tong University, Shanghai 200030, China; Department of Psychology (Scarborough), University of Toronto, Toronto, ON M1C 1A4, Canada; Department of Psychology, University of Oregon, Eugene, OR, 97403, USA

**Author notes:** Y. Zheng and D. Wang contributed equally as co-first authors. Correspondence: Sze Chai Kwok; Yao Li.

**Keywords:** Metacognition, DTI, superior longitudinal fasciculus, functional connectivity, structural integrity, precuneus

## Abstract

Metacognition as the capacity of monitoring one’s own cognition operates across domains. Here, we addressed whether metacognition in different cognitive domains rely on common or distinct neural substrates with combined diffusion tensor imaging (DTI) and functional magnetic resonance imaging (fMRI) techniques. After acquiring DTI and resting-state fMRI data, we asked participants to perform a temporal-order memory task and a perceptual discrimination task, followed by trial-specific confidence judgments. DTI analysis revealed that the structural integrity (indexed by fractional anisotropy) in the anterior portion of right superior longitudinal fasciculus (SLF) was associated with both perceptual and mnemonic metacognitive abilities. Using perturbed mnemonic metacognitive scores produced by inhibiting the precuneus using TMS, the mnemonic metacognition scores did not correlate with individuals’ SLF structural integrity anymore, revealing the relevance of this tract in memory metacognition. In order to further verify the involvement of several cortical regions connected by SLF, we took the TMS-targeted precuneus region as a seed in a functional connectivity analysis and found the functional connectivity between precuneus and two SLF-connected regions (inferior parietal cortex and precentral gyrus) differentially mediated mnemonic but not perceptual metacognition performance. These results illustrate the importance of SLF and a putative white-matter grey-matter circuitry that supports human metacognition.

## Introduction

The capacity of reflecting on one’s own cognitive process is known as metacognition (Flavell, 1979; Fleming & Dolan, 2012; Yeung & Summerfield, 2012). Given that metacognition has been considered as one of the most crucial functions emerged during evolution (Heyes, 2016), researchers have endeavored to understand how a metacognitive judgment is computed (Fleming & Daw, 2017; Kepecs et al., 2008; Zylberberg et al., 2016), how it is disrupted in psychiatric disorders (Hauser et al., 2017a; Rouault et al., 2018a), and how its accuracy can be improved (Carpenter et al., 2019). Metacognition is an umbrella term for the higher-level cognition about the lower-level cognition in various domains (e.g., perception and memory). An interesting and important question is whether the neural circuit supporting metacognition is the same or distinct across different cognitive domains (Rouault et al., 2018b).

A large body of functional magnetic resonance imaging (fMRI) work have demonstrated a nuanced picture for this domain-generality issue of metacognition. For example, the dorsolateral prefrontal cortex (DLPFC) might be involved in reading out the information of primary decision-making and using it for computing both perceptual and mnemonic metacognitive judgments (Chua & Ahmed, 2016; Fleming & Dolan, 2012; Kwok et al., 2019; Rounis et al., 2010). Morales et al. (2018) reported that the dorsal anterior cingulate cortex (dACC) was active during metacognitive evaluation in both memory and perception tasks but also found the activation patterns decoded from a perception task in the posterior medial frontal cortex and ventral medial prefrontal cortex predict metacognitive judgements in a memory task. In contrast to these domain-general components, other evidence also indicates domain-specific mechanisms. For example, accurate perceptual metacognition is dependent on the accessibility of performance-monitoring information coded in the dACC to the anterior PFC (Allen et al., 2017; Fleming et al., 2010, 2014; McCurdy et al., 2013), whereas accurate memorial metacognition is dependent on memory-mediated regions such as the medial PFC, mid/posterior cingulate cortex, inferior parietal lobule (IPL) and precuneus (Baird et al., 2013; Chua et al., 2006; McCurdy et al., 2013; Simons et al., 2010; Ye et al., 2018).

Although much progress has been made on how cortical networks support metacognition in different domains, less is known about how white matter pathways, through which interregional information communicates, contribute to supporting these cognitive processes. Fleming et al. (2010) reported that metacognitive ability on perception domain was positively correlated with the diffusion anisotropy in the genu of corpus callosum (the callosum forceps minor) which links the anterior PFC. To our knowledge, Baird et al. (2015) was the only study to date directly compared the white matter microstructure related to perceptual and mnemonic metacognitive ability. They found that accurate metacognitive evaluation on a perception task positively correlated with the diffusion anisotropy underlying the anterior cingulate cortex (ACC), whereas accurate metacognitive evaluation on a memory task positively correlated with the diffusion anisotropy of the white matter underlying the inferior parietal lobue (IPL), indicating metacognition in different cognitive domains might rely on distinct neural substrates. Nevertheless, Baird and colleagues did not control for the local task properties (visual stimuli vs verbal words) nor the metrics quantifying the perceptual and mnemonic metacognitive ability (2-AFC vs Y/N responses), which can bias the comparison across domains (Lee et al., 2018; Rouault et al., 2018b). They also did not characterize the diffusion property at specific, finely-defined locations along white matter tracts and their relationship with metacognitive abilities, so that the actual extent to which white matter tracts contributes to metacognition in each cognitive domain might be underestimated (Teubner-Rhodes et al., 2016).

Therefore, the present investigation sought to elucidate the neurobiological mechanisms underpinning metacognition across two different domains in relation to the white-matter diffusion property. We used a diffusion tensor imaging (DTI) tractography technique, named as automated fibre quantification (AFQ; Yeatman et al., 2012, 2014), to perform intra-tract analysis of tissue features along the neuronal fibre tracts. This method gives sensitive measures of white matter structural integrity and allows us to examine its relationship with metacognitive ability in different domains. Given the known neural substrates of metacognition, we selected the following five white matter tracts (two bilateral and one unilateral): The bilateral superior longitudinal fasciculus (SLF), which links the domain-general DLFPC to the precentral gyrus and inferior parietal cortex (Hecht et al., 2015); the bilateral cingulum bundle (CB), which connects the domain-general ACC to the posterior parietal regions (Heilbronner & Haber, 2014); and the callosum forceps minor (CFM), which connects the perceptual metacognition related regions such as the left and right anterior PFC (Fleming et al. 2010; Park et al., 2008).

In addition, making use of our published behavioral data (Ye et al., 2018), we further verified the relevance of mnemonic metacognition-related white matter tracts using a set of perturbed metacognition scores following TMS to the precuneus or to the vertex. We observed that the FA of the right SLF was significantly correlated with both perceptual and mnemonic metacognitive ability. However, with altered metacognitive scores induced by precuneus-TMS, the relationship of SLF FA with mnemonic metacognition disappeared, while the one with perceptual metacognition was preserved.

Finally, to fully elucidate how the afferent and efferent between precuneus and brain regions connected by the right SLF exclusively supports mnemonic metacognition, we ran a restingstate functional connectivity (rs-FC) analysis with the TMS-targeted precuneus site as the seed. The precuneus-right precentral gyrus and precunues-right IPL rs-FCs were found to be correlated with intact mnemonic metacognitive ability (but not the purtubed one); while these frontal-parietal network rs-FCs were not correlated with the perceptual metacognitive ability. Taken together, the current results indicate that the superior longitudinal fasciculus assumes an important anatomical function to support human metacognition.

## Materials and Methods

### Participants

Eighteen university students (7 females, aged 19-24 years) from East China Normal University participated in this study. All participants had normal or corrected-to-normal vision, reported no history of psychiatric and neurological diseases, and no other contraindications for MRI. All participants gave written informed consent, and were financially compensated for their participation. The study was approved by University Committee on Human Research Protection of East China Normal University. To guarantee the quality of metacognition estimation, we followed previous investigations (Allen et al., 2017; Morales et al., 2018; Rouault et al., 2018a), and excluded one participant whose first-order performance in the memory task was at chance level (d= −.04) and one participant who missed 10% of the trials in total (> 3SD from the average missed trials, which was 2.6% ± 2.2%). For each participant, we also discarded trials of which the first-order decision and confidence response time was faster than 100 ms or slower than three standard deviations from per-subject mean (2.1% of trials were discarded).

### Study design, Behavioral tasks, and Stimuli

The behavioral paradigm employed a two-way factorial design (2 Cognitive Tasks x 2 TMS Sites). In this study, participants were required to play a stimulus-matched memory (temporal order judgment; TOJ) task and a perception (visual discrimination) task. For the memory task, on day 1, participants played seven distinct chapters of a first-person perspective action-adventure video game *Beyond: Two Souls* with Sony PlayStation as the memory encoding period. Prior to each testing session, participants received 20-min repetitive TMS that targeted at the precuneus or the vertex (as control site) in a counterbalanced order (Figure 1A-B, see section *Repetitive TMS: procedure, protocol, and sites* below). In each trial of the TOJ memory task (Figure 1C), participants were presented with two images from the seven chapters of the video game they played, and were required to choose the one that occurred earlier. Images were presented for 5 s, followed by a 3-s confidence rating period where participants needed to report their confidence in the TOJ judgement. There were four available confidence ratings (i.e., “Very Low”, “Low”, “High”, or “Very High”), and participants were encouraged to make use of the whole confidence scale. After the confidence rating, participants entered the inter-trial interval with a variable duration from 1-6 s. In total, per subject, 240 pairs of images were extracted and paired up for each TOJ memory task session.

**Figure 1.**
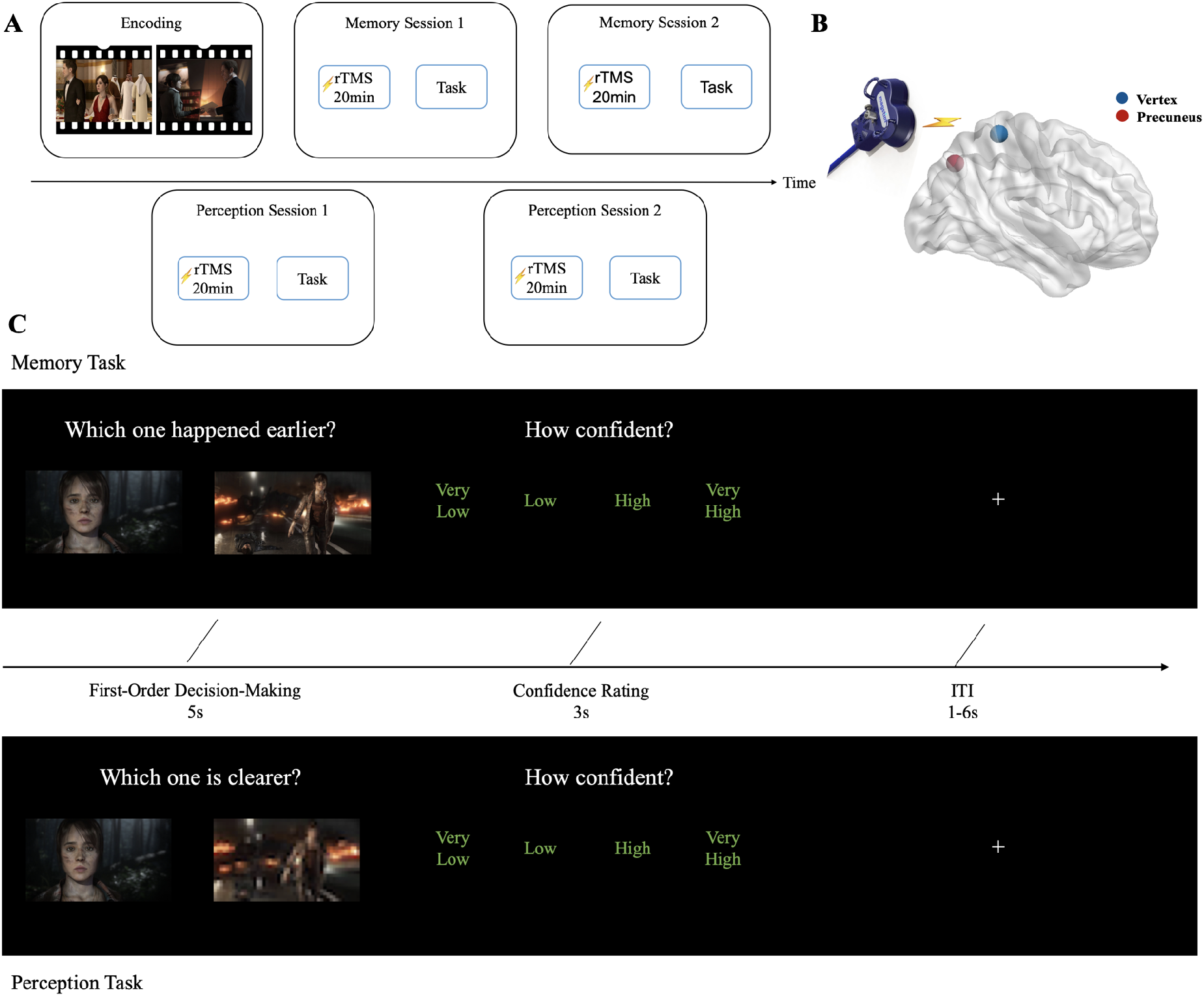
Study overview. ***A***, Experiment design. In order to alter participants’ metacognition scores, we applied 20-min rTMS to one of the two cortical sites prior to the main task in each session. ***B***, rTMS locations. The precuneus stimulation (MNI *x, y, z* = 6, −70, 44) was based on Kwok et al. (2012); vertex stimulation as a control site. ***C***, Task Procedure. In the memory (temporal order judgement) task, participants were required to choose the image that occurred earlier in the video game; in the perceptual (visual discrimination) task, participants were required to identify which image was clearer (or blurrier). In both tasks, after each first-order decision making, participants rated their confidence for the current trial.

The perception task followed a similar procedure with the memory task. In each trial, the same set of subject-specific paired images used in the memory task were presented to participants. However, we manipulated resolution differences between the images, and asked participants to report which one was clearer (or blurrier) and their associated confidence level (Figure 1C). The same sets of paired images in each memory task session were presented in each perception task session in an identical order, but with manipulation on the image resolutions. We used Python Imaging Library to reduce the resolution of one of the two images (i.e., resizing the image to change the pixel per inch [PPI]). The higher the PPI, the smaller the difference would be in the resolution between the resized and original images, thus the harder for participants to discriminate the clearer one. For example, if we reduced the PPI of one of the paired images to 30%, the image resolution difference would be 0.7; while if the PPI of one image was reduced to 0.9, the resolution difference would be 0.1. The range of the resolution difference for all image pairs was from 0.05 to 0.65. To match participants’ performance across the two tasks, we used an *n*-down/1-up adaptive staircase procedure to adjust the image resolution online and converge on ~71% performance.

There were 480 trials in total for each cognitive task (2 sessions × 4 blocks × 60 trials per block), and each task lasted around 45 min. A two-alternative forced choice (2-AFC) design was used for both the perception and memory task, and the experiment-related parameters (i.e., number of trials; dimension, position and sequence of the presented stimulus; time limits for responses; and the inter-trial intervals) were set identical in both tasks. All the visual stimuli were presented with E-prime software (Psychology Software Tools), and the presentation order of the paired images were counterbalanced throughout the experiment. To help participants get familiar with task demands, we had them do a practice block prior to each testing session. Stimuli for the practice blocks were not used in the main tests.

### MR Image Acquisition

High-resolution structural, DTI, and resting-state fMRI data were acquired on a separate day prior to the first session. All MR images were acquired using a 3.0 T Siemens Trio MRI scanner with a 32-channel head coil. High-resolution T1-weighted images were acquired using Magnetization Prepared Rapid Gradient Recalled Echo (MPRAGE) sequence with the following parameters: repetition time (TR) = 2530 ms, echo time (TE) = 2.34 ms, inversion time = 1100 ms, flip angle = 7°, field of view (FOV) = 256 mm, slice number = 192, voxel size = 1 × 1 × 1 mm^3^, slice thickness = 1.0 mm.

Seventy transverse DWI (diffusion-weighted imaging) slices were acquired with the parameters: TR = 11000 ms, TE = 98 ms, FOV = 256 mm, voxel size = 2 × 2 × 2 mm^3^, acquisition matrix = 128 × 128, phase encoding direction: anterior to posterior (A > > P), 60 gradient directions (b = 1000 s/mm^2^) and 2 non-diffusion images were obtained. The fMRI images were collected with the following parameters: TR = 2000 ms, TE = 30 ms, FOV = 230 mm, flip angle = 70°, voxel size = 3.6 × 3.6 × 4 mm^3^, slice number = 33, parallel to the AC-PC plane. For each subject, 220 whole-brain volumes were acquired.

### MR Image Processing

#### Tract-based analysis for DTI data

For tract-based analysis, we performed DTI image preprocessing using VISTASOFT package (http://web.stanford.edu/group/vista/cgi-bin/wiki/index.php/Software). After the correction of head motion and eddy-current distortion, the DWI images were registered to the averaged nondiffusion weighted images. After that, the DWI images were registered to the T1-weighted image and the corresponding fractional anisotropy (FA) maps were obtained. A higher FA value implies a stronger structural connectivity between the brain regions (Greicius et al., 2009). Fiber tracking was performed using Automating Fiber-Tract Quantification (AFQ, https://github.com/jyeatman/AFQ; Yeatman et al., 2012). First, a deterministic streamline tracking algorithm (Basser et al., 2000; Mori et al., 1999) was employed for whole-brain tractography. The tracking was performed within the white matter mask, seeded at the voxels with FA > 0.3 and terminated if the voxelwise FA value was below 0.2 or the minimum angle between last path segment and next step direction was larger than 30° (Yeatman et al., 2012). Second, we performed fiber tract segmentation using waypoint regions of interest (ROIs). The fiber was included in the fiber tract if it passes through two ROIs that define the trajectory of the fiber tract. The ROIs defined in Montreal Neurological Institute (MNI) space were registered to each participant’s native structural space through nonlinear transformation (Dougherty et al., 2005; Wakana et al., 2007). Finally, the candidate fibers were removed if passing through the white matter regions unlikely the parts in fiber tract probability map (Hua et al., 2008) or the three-dimensional Gaussian covariance of the sample points are larger than 5 s.d. from the mean (Yeatman et al., 2012). The obtained tract was centered in all the corresponding tract fibers. A curve was created by defining 100 evenly spaced sample points along each fiber and calculating the mean position of each sample point in the curve accordingly. The diffusion metric, FA, was calculated as a weighted average of each individual fiber’s measurement at each sample point. Five frontal-parietal white matter tracts (i.e., right/left superior longitudinal fasciculus, SLF; right/left cingulum bundle, CB; the callosum forceps minor, CFM) were selected for further analysis.

#### Functional connectivity analysis

For the FC calculation, the fMRI image processing was performed using Data Processing Assistant for Resting-State fMRI (DPARSF, https://www.nitrc.org/projects/dparsf/), which is based on Statistical Parametric Mapping 8 (SPM8, http://www.fil.ion.ucl.ac.uk/spm) and Resting-State fMRI Data Analysis Toolkit (REST v1.8, http://www.restfmri.net). The first 10 volumes were discarded for machine stabilization and participants’ adaption to the environment. The remaining 210 volumes were subsequently preprocessed by the following steps: slice-timing, head motion correction, normalization to the MNI space with voxel size resampled to 3 × 3 × 3 mm^3^, spatial smoothing using an isotropic Gaussian filter kernel with 6 mm full-width at half maximum (FWHM), linear detrending, nuisance signal regression and temporal band-pass filtering (0.01-0.10 Hz). No participant was excluded by the criterion that the head motion was above 3 mm or 3°. Nuisance regressors included Friston-24 head motion parameters (Friston et al., 1996), white matter and CSF signals and the global signal. The FC map was generated by computing the averaged time series within the 8-mm-radius sphere centered at the precuneus (MNI coordinate: *x* = 6, *y* = −70, *z* = 44; Kwok et al., 2012) and correlating them with the time series of all other grey matter voxels in the whole brain using Pearson’s correlation analyses. The FC map was transformed to Z-score map by Fisher’s Z transformation for statistical analysis.

#### MRI data statistical analysis

Pearson correlation analysis was performed between FCs or DTI metrics and cognitive scores (i.e. first-order performance, mean confidence level, and metacognitive efficiency) for all the subjects using IBM SPSS 20. For FCs, the correlations were corrected for multiple comparisons using AlphaSim (M = 1000,p < 0.001, cluster size = 13 voxels). The correlations between DTI metrics and cognitive cores were calculated using Pearson correlation analysis. For DTI metrics, the correlations were controlled for multiple-comparisons error with false discovery rate (FDR) correction at p < 0.05.

#### Behavioral data analysis

Type-1 memory and perceptual performance were quantified as d’, a signal detection theoretic measure of type-I sensitivity. Type-2 metacognitive sensitivity was estimated by meta-d’, which is expressed in the same scale as d’ and indicates the extent to which a participant could discriminate correct responses from incorrect ones. Here we used metacognitive efficiency (meta-d’ / d’) to represent participants’ metacognitive ability independent of primary decisionmaking performance. We quantified metacognitive ability through a hierarchical Bayesian Meta-d’ model (Fleming, 2017). This computational model gives rise to more precise meta-d’ estimation at both individual- and group-level (log-transformed) level, which allow direct and precise comparison and correlation analyses of metacognitive abilities across each subjects and each conditions. RStudio and IBM SPSS 22 were used for the behavioural data analysis.

#### Repetitive TMS: procedure, protocol, and sites

In order to verify the functional relevance of white-matter tracts and their functional connectivity in support of metacognition, we utilized our previously published behavioral data (Ye et al., 2018) for the DTI and functional connectivity analyses here. In that study, we used rTMS to alter subjects’ metacognitive scores. rTMS was applied using a Magstim Rapid^2^ magnetic stimulator connected to a 70 mm double air film coil (The Magstim Company, Ltd., Whitland, UK). To localize the target brain regions, Brainsight2.0 (Rogue Research Inc., Montreal, Canada) was used for the subject-specific structural T1-weighted images. Participants’ brains were normalised by transforming to the Montreal Neurological Institute (MNI) template. To prepare the subject-image registration and promote online processing of the neuronavigation system, four location information of each subject’s head were obtained manually by touching the tip of the nose, the nasion, and the inter-tragal notch of each ear using an infra-red pointer. In each session, rTMS was delivered to either the precuneus or vertex before the main task. TMS was applied at 1 Hz frequency for a continuous duration of 20 min (1200 pulses in total) at 110% of active motor threshold. The order of stimulation sites was counterbalanced across sessions. The coil was held to the scalp of the participant with a custom coil holder and the subject’s head was propped a comfortable position. Coil orientation was parallel to the midline with the handle pointing downward. The stimulation sites are in the precuneus with MNI coordinates *x* = 6, *y* = −70, *z*= 44 (Kwok et al., 2012) and in a control area on the vertex (Jung et al., 2016); the latter is identified at the point of the same distance to the left and the right pre-auricular, and of the same distance to the nasion and the inion (Figure 1B).

## Results

### Behavioral Results

By fitting the Bayesian hierarchical meta-d’ model, we observed that, in comparison to the vertex TMS, the precuneus TMS marginally significantly diminished the value of group-level metacognitive efficiency in the memory task (*p*_Δ*θ*>0_ ~ .92; 95% HDI = [−.068, 0.396]), but not in the perception task (*p*_Δ*θ*>0_ ~ .41; 95% HDI = [−.150, 0.120]; Figure 2B). The group-level metacognitive efficiency for memory and perception tasks were also significantly positively correlated in the vertex TMS condition (ρ=.526; 95% HDI = [.003, .964]). For the following neural analyses, we obtained the metacognitive efficiency scores in both memory and perception tasks for each single participant in both TMS conditions (Mean ± Standard Deviation: Memory-Vertex, 1.89±.50; Perception-Vertex, 1.31±.33; Memory-Precuneus, 1.68±.64; Perception-Precuneus 1.29±.20; Figure 2C]. These subject-level scores, including the ones in precunues TMS conditions, were used in the DTI and functional connectivity analyses.

**Figure 2.**
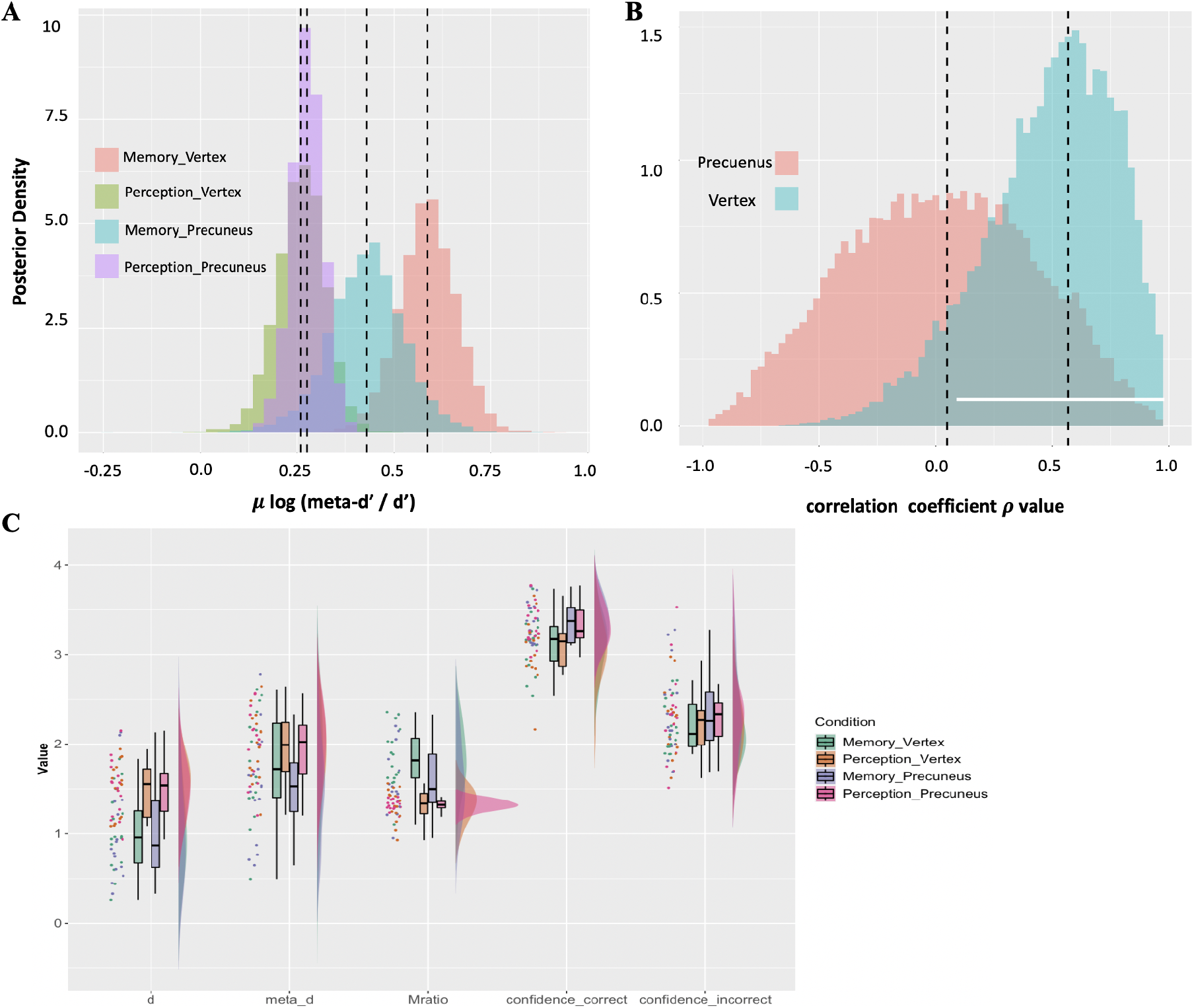
Behavioral results. ***A*.** Group posterior densities of metacognitive efficiency (log [meta-d’ / d’]) in perceptual and memory tasks in both TMS – precuneus and vertex conditions. ***B*.** Posterior densities of the correlation coefficient *ρ* between metacognitive efficiencies in perceptual and mnemonic domains in both TMS–vertex (control) condition. The white bar indicates the 95% Highest Density Interval (HDI) which excludes zero, indicating the correlation is significantly positive. The dotted lines in both figures show the ground-truth parameter values. ***C*.** Rain cloud plot for participants’ first-order performance, average confidence rating for both correct and incorrect trials, as well as their metacognitive sensitivity (i.e. meta-d’) and efficiency (i.e. Mratio) across different conditions. There is a significant main effect of d’ for cognitive domains (F(1,15)=77.263, p<.001, partial η^2^=.837 and significant main effects of cognitive domains and trial correctness on participants’ confidence ratings (Cognitive domains: F(1,15)=8.025, p=.013, partial η^2^ =.349; Trial Correctness: F(1,15)=243.190, p<.001, partial η^2^=.942). The participants performed better and had higher confidence in the perception task than in the memory task regardless of the TMS sites.

### Diffusion Tensor Imaging Results

Neurally, we report how FA in the white matter tracts might be correlated with participants’ metacognitive scores. First and foremost, we found a significant positive correlation between the FA in the anterior portion of right SLF and memory metacognition (nodes 1-53, FA: R = [0.56, 0.77], p_FDR_ < 0.05) as well as a significant positive correlation with their perceptual metacognition (nodes 1-40, FA: R = [0.59, 0.76], p_FDR_ < 0.05) in the TMS-vertex condition (Figure 3B-C, top panel). With TMS applied to the precuneus, the correlation between right SLF FA and perceptual metacognition are preserved (nodes 6-43, FA: R = [0.58, 0.74], p_FDR_ < 0.05; Figure 3C, bottom panel) wheres correlation between the SLF FA and memory metacognition did not survive FDR correction (Figure 3B, bottom panel).

**Figure 3.**
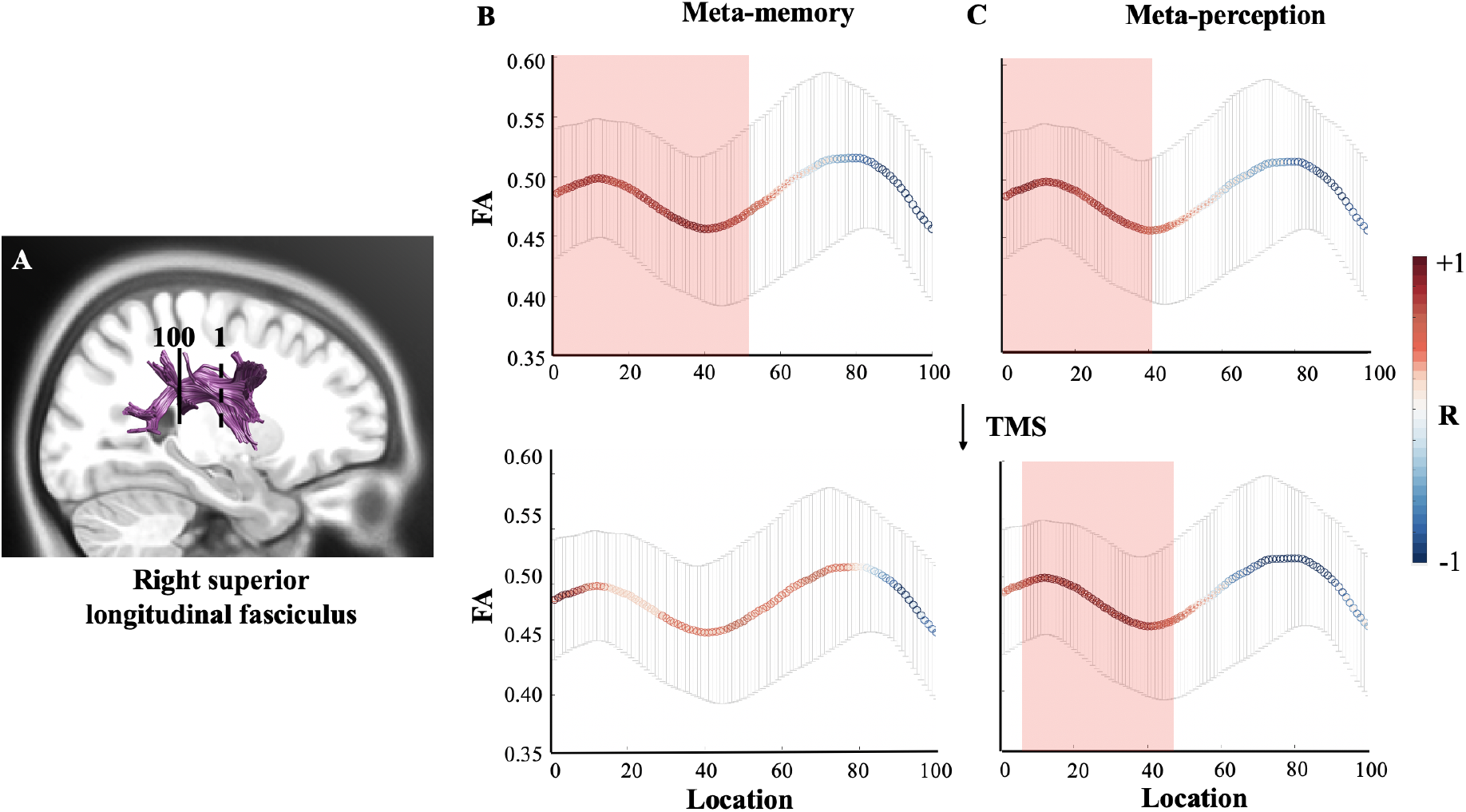
DTI results. The FAs of the nodes along the right anterior SLF had positive correlations with both perceptual and mnemonic metacognitive efficiency. ***A***, the portion of SLF that was tracked and evenly divided into 100 nodes by the AFQ method. The dash line represents the location of the starting point (marked by numeral 1) and the solid line represents the location of the ending point (marked by numeral 100). ***B*** and ***C*** respectively illustrates the correlation between the FA value of each SLF node and mnemonic or perceptual metacognitive efficiency (Top two panels: without TMS to precuneus; bottom two panels: TMS to precuneus). When precuneus-TMS diminished mnemonic metacognitive ability, the correlation between right SLF FA and memory metacognition disappeared. The x-axis is the individual node alongside the SLF, and its corresponding FA value is shown on y-axis. The colour of the curved lines illustrates the correlation between the FA of a SLF node and metacognitive efficiency scores. Those nodes that had significant correlation with metacognitive efficiency scores are marked by a red rectangular. The error bars denote standard error of the means over participants. Note: DTI, diffusion tensor imaging; FA, fractional anisotropy; AFQ, Automating Fiber-Tract Quantification; SLF, superior longitudinal fasciculus; TMS, transcranial magnetic stimulation.

This finding indicated that the right SLF play a domain-general role in supporting metacognition. To further examine this possibility, we modelled together both perceptual and mnemonic metacognitive efficiency scores when testing against the right SLF FA to see how the intact metacognitive ability in each domain (i.e., measured in the vertex TMS condition) might be uniquely correlated with the SLF structural integrity. A multiple linear regression analysis revealed that, after controling for covariance between the intact mnemonic and perceptual metacognitive effieceincy scores, neither of them significantly related to the FA of right SLF nodes (p_FDR_ > 0.05). This confirmed the importance of the right SLF in explaining the correlation between mnemonic and perceptual metacognition, suggesting domaingenerality of metacognition.

In order to provide further support for our claim that SLF FA is associated specifically with metacognition but not with other first-order processes, we ran the same analyses, now separately using either participants’ primary task accuracy (d’) or mean confidence ratings. We did not find any nodes along SLF that showed the diffusion properties significantly correlated with subjects’ first-order performance, nor with confidence ratings, in either of the cognitive tasks. These control tests indicate that the FA results cannot be explained by first-order performance or confidence, thus providing evidence that the putative effects are indeed metacognition-specific.

Moreover, we observed that the FA of the callosum forceps minor (CFM), which connects bilateral aPFC, exhibited positive correlations with mnemonic metacognition (nodes 9-13, 20-23, 92-95, FA: R = [0.50, 0.58], p_unc_ < 0.05) albeit at an uncorrected threshold. Similarly, the FA of the cingulum bundle portion, which extends to the dACC, also had positive correlations with both mnemonic and perceptual metacognitive ability (Mnemonic: nodes 28-34, 42-44, 86-92, FA: R = [0.51, 0.56], p_unc_ < 0.05; Perceptual: nodes 27-32, FA: R = [0.52, 0.64], p_unc_ < 0.05) at an uncorrected threshold. Interestingly, we did not replicate the CFM FA – perceptual metacognition relationship reported by Fleming and colleagues (2010) here, with possible reasons including the sub-domian differences within visual perception (Song et al., 2011). Since the results on CFM and cingulum bundle portion did not survive the correction threshold, we did not consider them further in the following functional connectivity analyses.

### Functional Connectivity MRI (fc-MRI) Results

Given that the mnemonic metacognitive efficiency scores did not correlate with the SLF FA any more after precunues TMS modulation, we selected the TMS-targeted precuneus as an anatomical seed and calculated its resting-state functional connectivity (rs-FC) with each voxel in the whole brain. This aimed to examine how the rs-FC predicted metacognitive abilities (measured in the TMS-control condition). For the memory domain, under the control TMS condition, the rs-FC between precuneus and two SLF-corssed clusters, the right caudal ventrolateral precentral gyrus (PrG; peak voxel MNI location, [42, −18, 3]) and the right IPL (peak voxel MNI location, [42, 27, 39]) (Figure 4A), were found to be significantly correlated with mnemonic metacognitive efficiency (Figure 4B). Notably, with precuneus-TMS disrupted mnemonic metacognitive efficiency scores, the relationships between these two rs-FCs and mnemonic metacognition both diminished. As expected, in contrast, no cluster survived correction when we repeated the same set of analyses when taking perceptual metacognitive efficiency into account in both TMS conditions. These FC results demonstrated the specificity of the information transimission between precuneus and right SLF-connected regions in supporting the mnemonic metacognitive ability alone, and further highlight the importance of the right SLF structural profile in mediating mnemonic metacognition.

**Figure 4.**
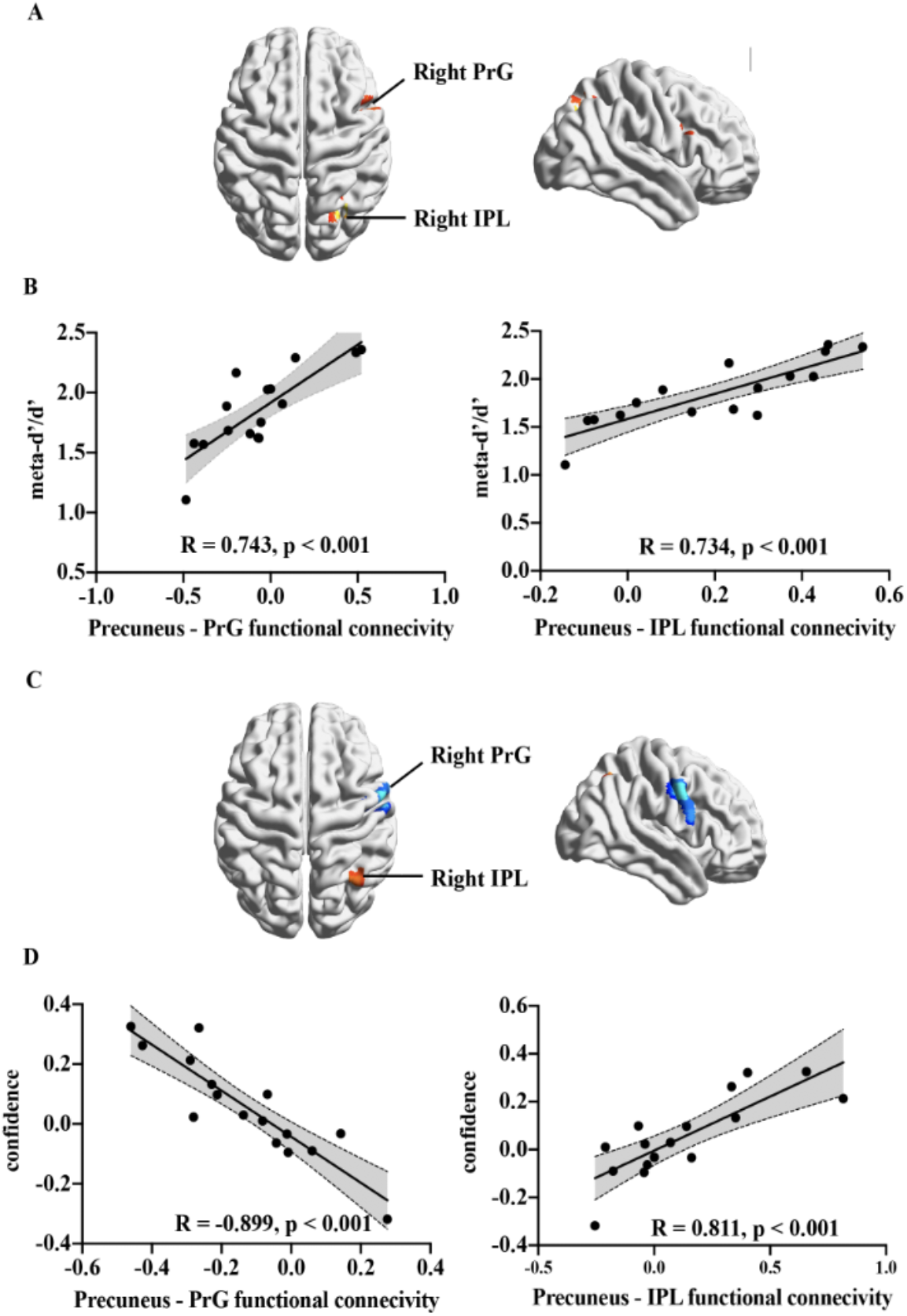
Functional connectivity results. With the TMS-targeted precuneus [6, −70, 44] selected as seed, ***A&B***, the Precuneus – right PrG FC and the Precuneus – right IPL FC both postively correlated with mnemonic metacognitive efficiency in the TMS-vertex condition. ***C&D***, the Precuneus – right PrG FC negatively correlated with the overall confidence in the memory task in the TMS-vertex condition, wheras the Precuneus – right IPL FC postively correlated with the overall confidence in this condition.

For control analyses, instead of metacognitive efficiency scores, we entered individual subjects’ first-order task performance and confidence ratings to repeat the same sets of FC analyses. Under the TMS-vertex condition, the clusters of PrG and IPL did not appear when modelling with the first-order memory performance. Interestingly, in terms of confidence rating, the precuneus – right IPL rs-FC positively correlated with the confidence in the memory task (peak voxel MNI location, [36, −57, 54], r=.811 p<.001), whereas the precuneus – right PrG rs-FC negatively correlated with the confidence in the task (peak voxel MNI location, [57, 0, 39], r=−.899, p<.001; Figure 4C,D). In TMS-precuneus condition, no significant correlations were observed in any of these tests.

## Discussion

Whether the computation of metacognition relies on the same mechanism across cognitive domains has been a controversial issue. In the present study, we addressed this domain-generality/specificity issue at the white matter structural integrity and grey matter functional connectivity level. By using DTI and resting-state fMRI techniques, we found metacognitive performance in perception and memory tasks are related to the superior longitudinal fasciculus and its connected regions.

We observed that the right SLF serves a domain-general role on metacognition. The structural integrity of the anterior portion of this tract correlates significantly with metacognitive efficiency in both perception and memory tasks, but not with primary decision-making performance nor confidence rating level. This specific portion of the right SLF underlies the precentral gyrus and links the right DLPFC to the right IPL (Hecht et al., 2015). Additionally, it also lies within the frontal lobe, which is not mixed by the temporal fibres in the arcuate fasciculus (Wakana, 2007) and is indicative of the DLPFC – IPL communication (Yeatman et al., 2012). On the one end, the right DLPFC was involved in both perceptual and mnemonic metacognition in both humans and monkeys, and suggested as reading out information related to primary decision-making and computing metacognitive judgements (Fleming and Dolan, 2012; Kwok et al., 2019; Morales et al., 2018; Shekhar and Rahnev, 2018). On the other end, the right IPL has been identified as crucial in mnemonic metacognition (Berryhil et al., 2007; Davidson et al., 2008; Simons et al., 2010). The right IPL has been reported as the “output gating” of working memory for information selection (Baddeley, 2000; Vilberg and Rugg, 2008; Wallis et al., 2015). Some studies reported that individuals would selectively ignore evidence favouring the unchosen alternatives during metacognitive evaluation (Aitchison et al., 2015; Samaha et al., 2016; Zylberberg et al., 2012), which implies the importance of such working-memory gating mechanism in metacognition. Thus, given the functional meaning of white-matter structural integrity, we propose that, in a metacognitive process, the right IPL select what information regarding the previous decision-making should be used and convey it to the right DLPFC through SLF for further computation. The better structural integrity of the right SLF is, the lower noise will be added during the information transmission, allowing the right DLPFC to read out information more precisely for the metacognitive computation.

The causal role of the right SLF to metacognition was further confirmed with TMS modulation on the precuneus. Applying TMS on the precunues, we disrupted participants’ mnemonic metacognitive ability, and observed that the SLF FA – mnemonic metacognition relationship did not exist anymore (while the SLF FA – perceptual metacognition remained). To unpack this TMS effect and investigate how the precuneus communicates with SLF-connected regions to support metacognition, we chose the targeted precuneus region as seed and examined how its resting-state functional connectivity correlated with the metacognitive efficiency scores (Hampson et al., 2006; Seeley et al., 2007). A frontal-parietal network (precuneus – IPL and precuneus – PrG) was found exclusively related to the mnemonic metacognition under the TMS control condition. The functional relevance of this network disappeared when TMS was applied at the precuneus. This result demonstrated that, although the structural integrity of right SLF supports both perceptual and mnemonic metacognition, the information afferent and efferent along the SLF could be domain-specific.

In the literature, the precuneus has been linked to generating mental images to aid detailed episodic memory retrieval (Fletcher et al., 1995; Hebscher et al., 2020; Koch et al., 2018; Richter et al., 2016; Sreekumar et al., 2018), as well as realizing temporal organization of episodic details (Foudil et al., 2020; Hasson et al., 2015; Kwok et al., 2014). In a recent investigation, Robin et al. (2015) reported a frontal-parietal network including the right precuneus and PrG was activated during episodic memory retrieval. We also observed evidence that the precuneus – right IPL and precuneus – right PrG communication support mnemonic metacognition via modulating the confidence level. The IPL might be involved in the information sampling and selection processes (e.g., Baddeley, 2000), and thus the more first-order memory-related inputs received from the precuneus, the higher and more accurate confidence rating would be. The PrG or a broader presupplmentary motor area (pre-SMA) region, which interconnects with the DLPFC and ACC, might be involved in cognitive control processes (Morales et al., 2018; Völker et al., 2018), and thus when conflicting signals are detected, this region might access more first-order memory-related information from the precuneus to guide a more accurate confidence rating. Therefore, our findings on the relationship of the right IPL – precuneus and right PrG – precuneus rs-FCs with mnemonic metacognitive ability lend support to the hypothesis that mnemonic metacognition relies on the read out of memory trace (Nelson & Narens, 1990). Together with the precuneus – hippocampus functional link (Ye et al., 2019), a cortico-hippocampal network of meta-mnemonic process is at work with primary mnemonic processes (McClelland et al., 1995; Wang et al., 2014; Zeidman et al., 2015), implicating the hippocampus in initiating the retrieval of the memory traces, which are then integrated to detailed recollections and conscious monitoring supported by the precuneus and the IPL (Richter et al., 2016).

Given that the SLF FA was not related to primary TOJ performance, in line with previous investigations (Allen et al., 2017; Bang et al., 2019; Fleming et al., 2014; Hauser et al., 2017b; Qiu et al., 2018; Rounis et al., 2010; Shekhar & Rahnev, 2018), our results could also lend support to the hierarchical model of metacognition (Fleming & Daw, 2017), which proposes that the metacognition evaluation is an inference based on primary decision-making evidence and other sources of information. The functional relevance of SLF might further provide evidence that how the domain-general hierarchical inference is implemented in the brain.

On a more speculative note, metacognition has been associated with states of consciousness (Lau & Rosenthal, 2011), and our current results might help facilitate the understanding on the neurocognitive mechanisms underlying consciousness. Several studies have revealed that subjective conscious awareness is dissociated from the objective visual perception, and requires the involvement of the prefrontal and parietal cortices apart from primary sensory regions (Colás et al., 2019; Del Cul et al., 2009; Lau & Passingham, 2006; Persaud et al., 2011). It has also been argued that to generate conscious experiences, a higher-level neural circuit needs to metacognitively access the primary sensory representation from the lower-level neural circuit to the working memory (Dehanene & Changeux, 2011; Lau & Rosenthal, 2011; Shea & Frith, 2019). Indeed, our results imply that, to form a conscious metacognitive evaluation, the IPL firstly represents first-order decision-making related information in the memory and then conveys it to the DLPFC via the SLF for further monitoring. Incidentally, in the patients with right DLPFC lesion the structural integrity of the right SLF was significantly correlated with their subjective visual conscious experiences but not objective perception task performance (Colás et al., 2019). Moreover, the white-matter volume of the right SLF also predicts the metacognitive beliefs of schizophrenia patients (Spalletta et al. 2014). These clinical findings are in line with ours in the health, thereby reinforcing the possibility that the SLF is indeed needed for the higher-level conscious assessment of mental processes.

There are several caveats to consider. First, the current study only concerns the two broad cognitive domains of perception and memory. Future research could test whether our “domain-general” results are applicable to other cognitive domains. Second, further research could employ the diffusion spectrum imaging (DSI) technique to better examine the structural integrity of the SLF and its functional relevance with metacognition by excluding the effects of crossing fibres. Future research could also consider using task-state fMRI together with dynamic casual modelling (Friston, 2011) to draw a more comprehensive picture of the neural circuit of metacognition. Particularly, given the TMS might also have an effect on remote brain regions (Bestmann et al., 2005), future research indeed worth combining the task-state BOLD signal to further understand how the TMS on the precuneus influences the mnemonic metacognition-related information transmission via the right SLF.

In conclusion, our findings reveal how the structural integrity of the right SLF supports both perceptual and mnemonic metacognition, and how the efferent and afferent between the precuneus and the right SLF-connected regions exclusively support mnemonic metacognition. The results reinforce the hypothesis that human metacognition relies on the superior longitudinal fasciculus to support metacognitive computation for both domain-general and domain-specific processes.

